# Combining citizen science and deep learning to amplify expertise in neuroimaging

**DOI:** 10.1101/363382

**Authors:** Anisha Keshavan, Jason D. Yeatman, Ariel Rokem

**Affiliations:** University of Washington; Institute for Neuroengineering; eScience Institute; Department of Speech and Hearing

## Abstract

Research in many fields has become increasingly reliant on large and complex datasets. “Big Data” holds untold promise to rapidly advance science by tackling new questions that cannot be answered with smaller datasets. While powerful, research with Big Data poses unique challenges, as many standard lab protocols rely on experts examining each one of the samples. This is not feasible for large-scale datasets because manual approaches are time-consuming and hence difficult to scale. Meanwhile, automated approaches lack the accuracy of examination by highly trained scientists and this may introduce major errors, sources of noise, and unforeseen biases into these large and complex datasets. Our proposed solution is to 1) start with a small, expertly labelled dataset, 2) amplify labels through web-based tools that engage citizen scientists, and 3) train machine learning on amplified labels to emulate expert decision making. As a proof of concept, we developed a system to quality control a large dataset of three-dimensional magnetic resonance images (MRI) of human brains. An initial dataset of 200 brain images labeled by experts were amplified by citizen scientists to label 722 brains, with over 80,000 ratings done through a simple web interface. A deep learning algorithm was then trained to predict data quality, based on a combination of the citizen scientist labels that accounts for differences in the quality of classification by different citizen scientists. In an ROC analysis (on left out test data), the deep learning network performed as well as a state-of-the-art, specialized algorithm (MRIQC) for quality control of T1-weighted images, each with an area under the curve of 0.99. Finally, as a specific practical application of the method, we explore how brain image quality relates to the replicability of a well established relationship between brain volume and age over development. Combining citizen science and deep learning can generalize and scale expert decision making; this is particularly important in emerging disciplines where specialized, automated tools do not already exist.

## Introduction

Many research fields ranging from astronomy, to genomics, to neuroscience are entering an era of Big Data. Large and complex datasets promise to address many scientific questions, but they also present a new set of challenges. For example, over the last few years human neuroscience has evolved into a Big Data field. In the past, individual groups would each collect their own samples of data from a relatively small group of individuals. More recently, large data sets collected from many thousands of individuals are increasingly more common. This transition has been facilitated through assembly of large aggregated datasets, containing measurements from many individuals, and collected through consortium efforts such as the Human Connectome Project (Glasser et al., 2016). These efforts, and the large datasets that they are assembling, promise to enhance our understanding of the relationship between brain anatomy, brain activity and cognition. The field is experiencing a paradigm shift (Fan et al., 2014), where our once established scientific procedures are morphing as dictated by the new challenges posed by large datasets. We’ve seen a shift from desktop computers to cyberinfrastructure (Van Horn and Toga, 2013), from small studies siloed in individual labs to an explosion of data sharing initiatives (Ferguson et al., 2014; Poldrack and Gorgolewski, 2014), from idiosyncratic data organization and analysis scripts to standardized file structures and workflows (Gorgolewski et al., 2016, 2017b), and an overall shift in statistical thinking and computational methods (Fan et al., 2014) that can accommodate large datasets. But one often overlooked aspect of our protocols in neuroimaging has not yet evolved to the needs of Big Data: expert decision making.

Specifically, decisions made by scientists with expertise in neuroanatomy and MRI methods (i.e., neuroimaging experts) through visual inspection of imaging data cannot be accurately scaled to large datasets. For example, when inspecting an MRI image of the brain, there is extensive variation in neuroanatomy across individuals, and variation in image acquisition and imaging artifacts; knowing which of these variations are acceptable versus abnormal comes with years of training and experience. Specific research questions require even more training and domain expertise in a particular method, such as tracing anatomical regions of interest (ROIs), editing fascicle models from streamline tractography (Jordan et al., 2017a), evaluating cross-modality image alignment, and quality control of images at each stage of image processing. On large datasets, especially longitudinal multisite consortium studies, these expert decisions cannot be reliably replicated because the timeframe of these studies is long, individual experts get fatigued, and training teams of experts is time consuming, difficult and costly. As datasets grow to hundreds of thousands of brains it is no longer feasible to depend on manual interventions.

One solution to this problem is to train machines to emulate expert decisions. However, there are many cases in which automated algorithms exist, but expert decision-making is still required for optimal results. For example, a variety of image segmentation algorithms have been developed to replace manual ROI editing, with Freesurfer (Fischl, 2012), FSL (Patenaude et al., 2011), ANTS (Avants et al., 2011), and SPM (Ashburner and Friston, 2005) all offering automated segmentation tools for standard brain structures. But these algorithms were developed on a specific type of image (T1-weighted) and on a specific type of brain (those of healthy controls). Pathological brains, or those of children or the elderly may violate the assumptions of these algorithms, and their outputs often still require manual expert editing. Similarly, in tractography, a set of anatomical ROIs can be used to target or constrain streamlines to automatically extract fascicles of interest (Catani and Thiebautdeschotten, 2008; Yeatman et al., 2012). But again, abnormal brain morphology resulting from pathology would still require expert editing (Jordan et al., 2017b). The delineation of retinotopic maps in visual cortex is another task that has been recently automated (Benson et al., 2014, 2012), but these procedures are limited to only a few of the known retinotopic maps and substantial expertise is still required to delineate the other known maps (Winawer and Witthoft, 2017; Wandell and Winawer, 2011). Another fundamental step in brain image processing that still requires expert examination is quality control. There are several automated methods to quantify image quality, based on MRI physics and the statistical properties of images, and these methods have been collected under one umbrella in an algorithm called MRIQC (Esteban et al., 2017). However, these methods are specific to T1-weighted images, and cannot generalize to different image acquisition methods. To address all of these cases, and scale to new, unforeseen challenges, we need a general-purpose framework that can train machines to emulate experts for any purpose, allowing scientists to fully realize the potential of Big Data.

One general solution that is rapidly gaining traction is deep learning. Specifically, convolutional neural networks (CNNs) have shown promise in a variety of biomedical image processing tasks. Modeled loosely on the human visual system, CNNs can be trained for a variety of image classification and segmentation tasks using the same architecture. For example, the U-Net (Ronneberger et al., 2015) which was originally built for segmentation of neurons in electron microscope images, has also been adapted to segment macular edema in optical coherence tomography images (Lee et al., 2017b), to segment breast and fibroglandular tissue (Dalmiş et al., 2017), and a 3D adaptation was developed to segment the Xenopus kidney (Çiçek et al., 2016). Transfer learning is another broadly applicable deep learning technique, where a number of layers from pretrained network are retrained for a different use case. This can drastically cut down the training time and labelled dataset size needed (Ahmed et al., 2008; Pan and Yang, 2010). For example, the same transfer learning approach was used for brain MRI tissue segmentation (gray matter, white matter, and CSF) and for multiple sclerosis lesion segmentation (Van Opbroek et al., 2015). Yet despite these advances in deep learning, there is one major constraint to generalizing these methods to new imaging problems: a large amount of labelled data is still required to train CNNs. Thus, even with the cutting-edge machine learning methods available, researchers seeking to automate these processes are still confronted with the original problem: how does a single expert create an annotated dataset that is large enough to train an algorithm to automate their expertise through machine learning?

We propose that citizen scientists are a solution. Specifically, we hypothesize that citizen scientists can learn from, and amplify expert decisions, to the extent where deep learning approaches become feasible. Rather than labelling hundreds or thousands of training images, an expert can employ citizen scientists to help with this task, and machine learning can identify which citizen scientists provide expert-quality data. As a proof of concept, we apply this approach to brain MRI quality control (QC): a binary classification task where images are labelled “pass” or “fail” based on image quality. QC is a paradigmatic example of the problem of scaling expertise, because a large degree of subjectivity still remains in QC. Each researcher has their own standards as to which images pass or fail on inspection, and this variability may have problematic effects on downstream analyses, especially statistical inference. Effect size estimates may depend on the input data to a statistical model. Varying QC criteria will add more uncertainty to these estimates, and might result in replication failures. For example, in (Ducharme et al., 2016b), the authors found that QC had a significant impact on their estimates of the trajectory of cortical thickness during development. They concluded that post-processing QC (in the form of expert visual inspection) is crucial for such studies, especially due to motion artifacts in younger children. While this was feasible in their study of 398 subjects, this would not be possible for larger scale studies like the ABCD study, which aims to collect data on 10,000 subjects longitudinally (Casey et al., 2018). It is therefore essential that we develop systems that can accurately emulate expert decisions, and that these systems are made openly available for the scientific community.

To demonstrate how citizen science and deep learning can be combined to amplify expertise in neuroimaging, we developed a citizen-science amplification and CNN procedure for the openly available Healthy Brain Network dataset (HBN; (Alexander et al., 2017)). The HBN initiative aims to collect and publicly release data on 10,000 children over the next 6 years to facilitate the study of brain development and mental health through transdiagnostic research. The rich dataset includes MRI brain scans, EEG and eye tracking recordings, extensive behavioral testing, genetic sampling, and voice and actigraphy recordings. In order to understand the relationship between brain structure (based on MRI) and behavior (EEG, eye tracking, voice, actigraphy, behavioral data), or the association between genetics and brain structure, researchers require high quality MRI data.

In this study, we crowd-amplify image quality ratings and train a CNN on the first and second data releases of the HBN (n=722), which can be used to infer data quality on future data releases. We also demonstrate how choice of QC threshold is related to the effect size estimate on the established association between age and brain tissue volumes during development (Lebel and Beaulieu, 2011). Finally, we show that our approach of deep learning trained on a crowd-amplified dataset matches state-of-the-art software built specifically for image QC (Esteban et al., 2017). We conclude that this novel method of crowd-amplification has broad applicability across scientific domains where manual inspection by experts is still the gold-standard.

## Results

### Overview

Our primary goals were to 1) amplify a small, expertly labelled dataset through citizen science, 2) train a model that optimally combines citizen scientist ratings to emulate an expert, 3) train a CNN on the amplified labels, and 4) evaluate its performance on a validation dataset. Figure 1 shows an overview of the procedure and provides a summary of our results. At the outset, a group of neuroimaging experts created a gold-standard quality control dataset on a small subset of the data (n=200), through extensive visual examination of the full 3D volumes of the data. In parallel, citizen scientists were asked to “pass” or “fail” two-dimensional axial slices from the full dataset (n=722) through a web application called braindr that could be accessed through a desktop, tablet or mobile phone (https://braindr.us). Amplified labels, that range from 0 (fail) to 1 (pass), were generated from citizen scientist ratings. A receiver operating characteristic (ROC) curve was generated for both the ratings averaged across citizen scientists and labels generated by fitting a classifier that weights ratings more heavily for citizen scientists who more closely matched the experts in the subset rated by both (gold-standard). Next, a neural network was trained to predict the weighted labels. The AUC for the predicted labels on a left out dataset was 0.99.

**Figure 1:**
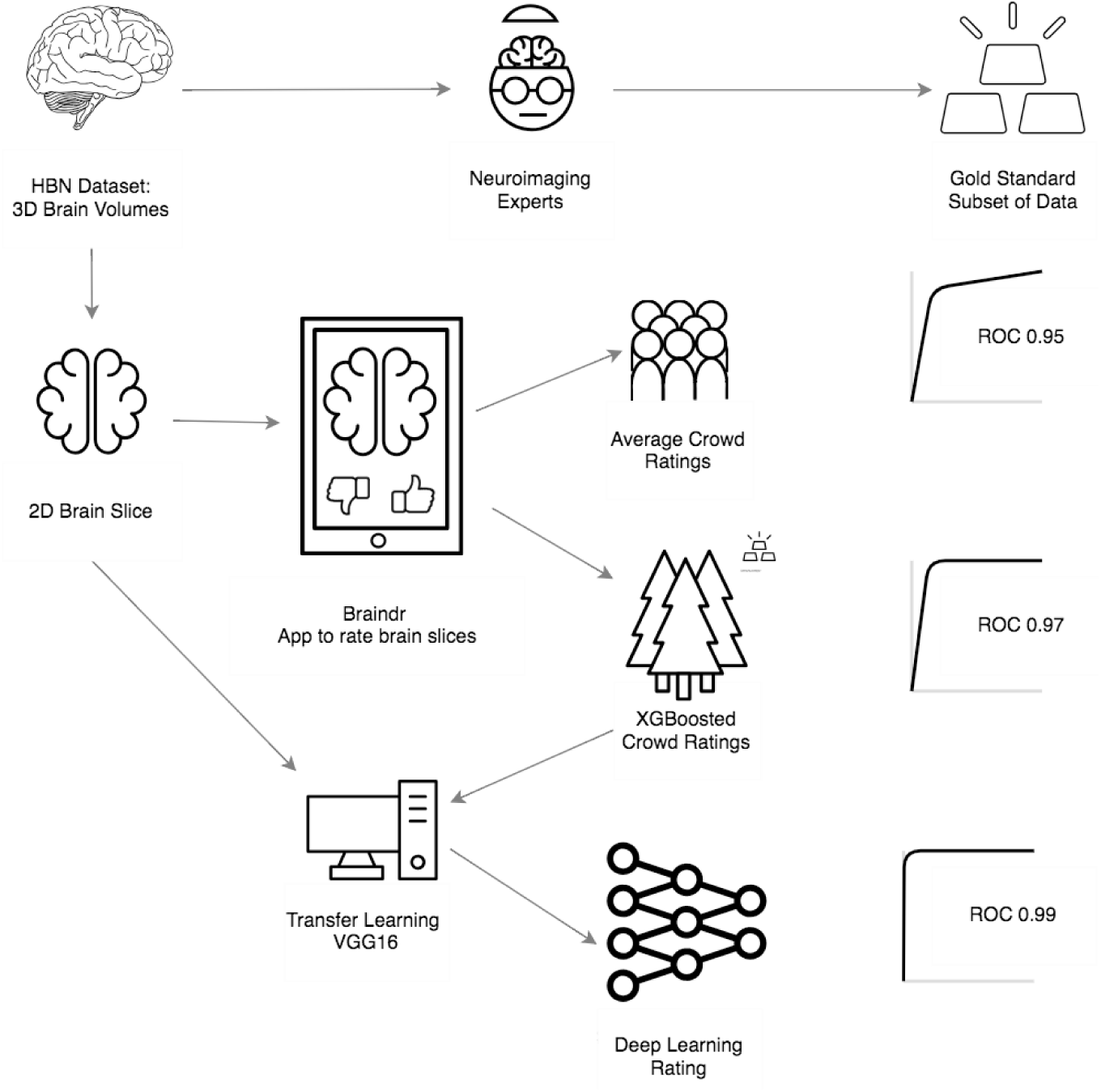
Overview and results of our procedure: First, the HBN data set was rated by 4 neuroimaging experts to create a gold standard subset of data. Next, the 3D MRI scans were converted into 2D axial brain slices, which were loaded onto braindr (https://braindr.us), a web application to crowdsource the quality ratings (see Methods). Area under the curve of a the Receiver Operating Characteristic curve (AUC) was calculated for the average citizen scientist quality rating for each slice. Compared to an expert-labeled test set, this resulted in an AUC of 0.95. In an effort to remove unreliable citizen scientists, the ratings were aggregated by fitting a model that weights each citizen scientist contribution to the slice score by how much that individual’s scores match those of the experts. The resulting AUC was 0.97. Finally, the 2D brain slices together with the weighted citizen scientist ratings were used to train a neural network. In an ROC analysis on left out data, the AUC of these predictions was 0.99.

### Aggregating Citizen Scientist Ratings to Emulate Expert Labels

Citizen scientists who rated images through the braindr web application differed substantially in terms of how well their ratings matched the experts’ ratings on the gold-standard subset: while some provided high-quality ratings that agree with the experts most of the time, others displayed variable and unreliable ratings. In order to capitalize on citizen scientists to amplify expert ratings to new data, a weighting of each citizen scientist was learned based on a reliable match to expert agreement in slices from the gold-standard set. We used the XGBoost algorithm (Chen and Guestrin, 2016a), an ensemble method that combines a set of weak learners (decision trees) to fit the gold-standard labels based on a set of features. In our case, the features were the average rating of the slice image from each citizen scientist (some images were viewed and rated more than once, so image ratings could vary between 1=always “pass” and 0=always “fail”). We then used the weights to combine the ratings of the citizen scientists and predict the left out test set. Figure 2A shows ROC curves of classification on the left-out test set for different training set sizes, compared to the ROC curve of a baseline model in which equal weights were assigned to each citizen scientist. We see an improvement in the AUC of the XGBoosted labels (0.97) compared to the AUC of the equi-weighted labels (0.95). Using the model trained on two-thirds of the gold standard data (n=670 slices), we extracted the probability scores of the classifier on all slices (see Figure 2B). The distribution of probability scores in Figure 2B matches our expectations of the data; a bimodal distribution with peaks at 0 and 1, reflecting that images are mostly perceived as “passing” or “failing”. The XGBoost model also calculates a feature importance score (F). F is the number of times that a feature (in our case, an individual citizen scientist) has split the branches of a tree, summed over all boosted trees. Figure 2C shows the feature importance for each citizen scientist, and 2D shows the relationship between a citizen scientist’s importance compared to the number of images they rated. In general, the more images a citizen scientist rates, the more important they are to the model. However, there are still exceptions where a citizen scientist rated many images and their ratings were incorrect or unreliable, so the model gave them less weight during aggregation.

**Figure 2:**
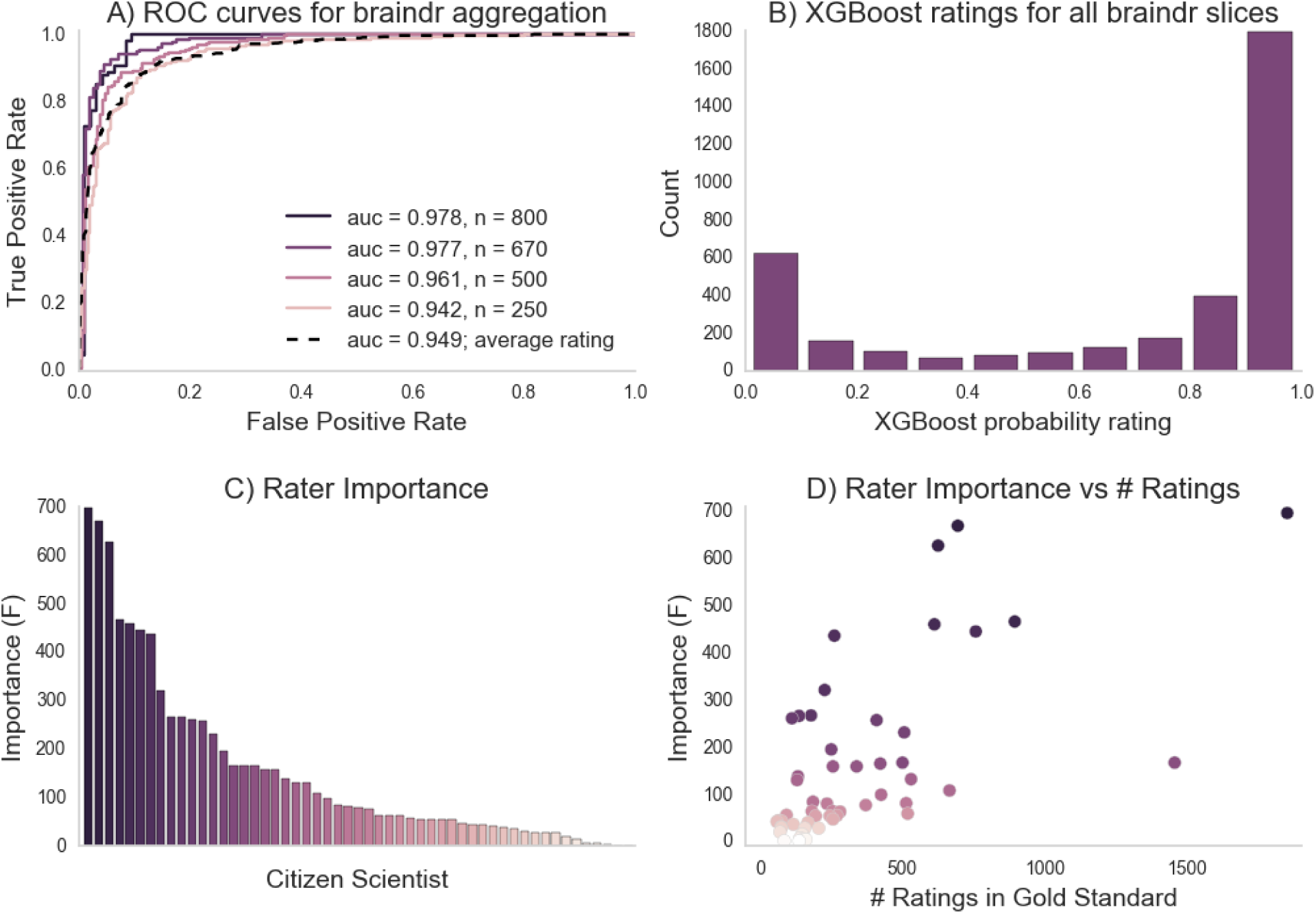
Braindr rating aggregation and citizen scientist importance: A. ROC curves on the test set for various training set sizes (here n denotes the number of training slices used). The dashed line is the ROC curve of the average citizen scientist ratings for all slices. B. The distribution of XGBoost probability scores on all Braindr slices. C. Feature importance for each anonymized user. D. Relationship between citizen scientist importance and total number of ratings in the gold-standard dataset.

### Training Deep Learning to Automate Image Labeling

Citizen scientists accurately amplify expert ratings but, ideally, we would have a fully automated approach that can be applied to new data as it becomes available. Thus, we trained a deep learning model to predict the XGBoosted labels that were based on aggregated citizen scientist ratings. A VGG16 neural network (Simonyan and Zisserman, 2014) pretrained on the ImageNet challenge dataset (Russakovsky et al., 2015) was used: we removed the top layer of the network, and then trained a final fully-connected layer followed by a single node output layer. The training of the final layer was run for 50 epochs and the best model on the validation set was saved. To estimate the variability of training, the model was separately trained through 10 different training courses, each time with a different random initialization seed. Typically, training and validation loss scores were equal at around 10 epochs, after which the model usually began to overfit (training error decreased, while validation error increased, see Figure 3A). In each of the 10 training courses, we used the model with the lowest validation error for inference on the held out test set, and calculated the ROC AUC. AUC may be a problematic statistic when the test-set is imbalanced (Saito and Rehmsmeier, 2015), but in this case, the test-set is almost perfectly balanced (see Methods). Thus, we found that a deep learning network trained on citizen scientist generated labels was a better match to expert ratings than citizen scientist generated labels alone: the deep learning model had an AUC of 0.99 (+/- standard deviation of 0.12, see Figure 3B).

**Figure 3:**
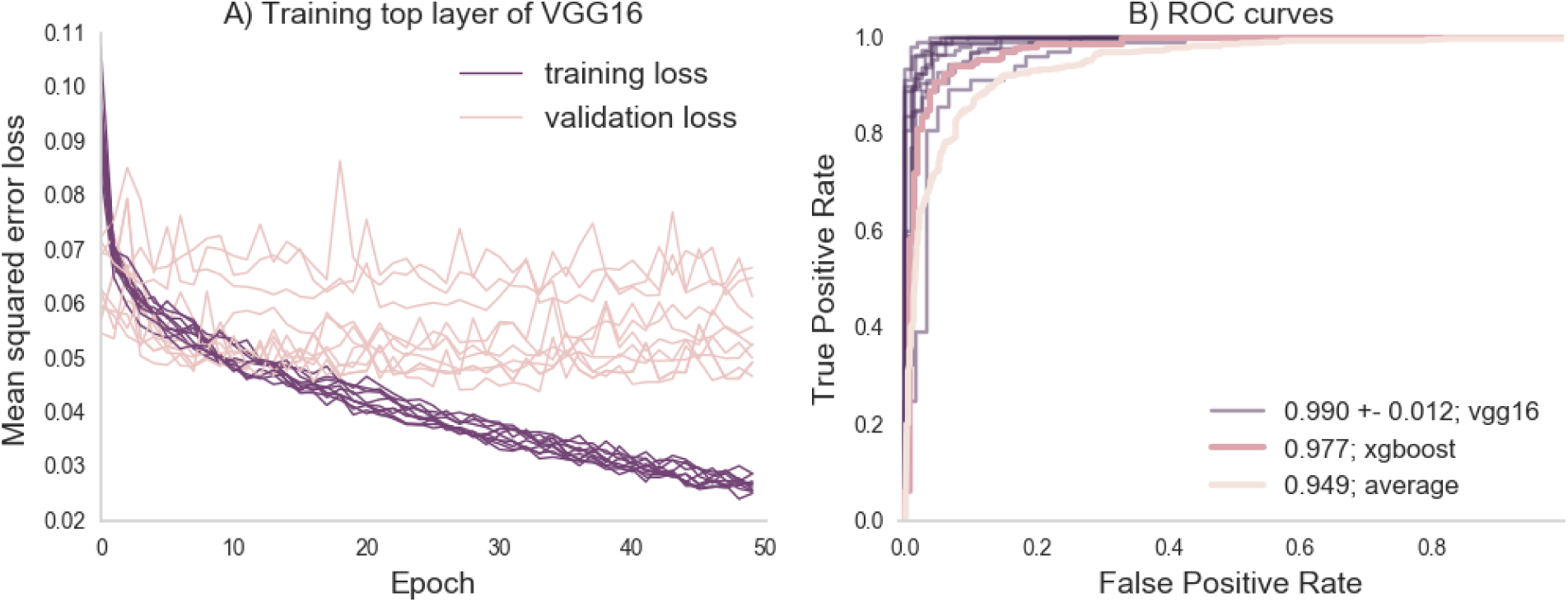
Deep learning training and evaluation on the left out test set: Part A shows the training and validation loss scores for 10 training runs, each with a different initialization seed. The training loss tends towards 0 but the validation loss plateaus between 0.05 and 0.07 mean squared error at the 10th epoch. Part B shows the ROC curve of the prediction on the test set against the binary classified gold-standard slices, along with the ROC curves computed from previous analysis (the average citizen scientist rating, and the XGBoosted ratings).

### Crowd amplification and deep learning strategy performs as well as a specialized QC algorithm

We validated our generalized approach of crowd-amplification and deep learning by comparing classification results against an existing, specialized algorithm for QC of T1 weighted images, called MRIQC (Esteban et al., 2017). The features extracted by MRIQC are guided by the physics of MR image acquisition and by the statistical properties of images. An XGBoost model was trained on the features extracted by MRIQC on a training subset of gold-standard images, and evaluated on a previously unseen test subset. The AUC was also 0.99, matching the performance of our crowd-trained deep learning model.

### Braindr-based quality control has a substantial impact on effect size estimates

The secondary goal of this study was to investigate how scaling expertise through citizen science amplification affects scientific inferences from these data. For this proof of concept, we studied brain development, which is the primary focus on the HBN dataset. Lebel and colleagues (Lebel and Beaulieu, 2011) found that increases in white matter volume and decreases in gray matter volume are roughly equal in magnitude, resulting in no overall brain volume change over development in late childhood. Based on Figure 2 in the Lebel manuscript (Lebel and Beaulieu, 2011), we estimate an effect of approximately −4.3 cm^3^ per year - a decrease in gray matter volume over the ages measured (See Figure 2 in the the original manuscript; we estimate the high point to be 710 cm^3^ and the low point to be 580 cm^3^ with a range of ages of approximately 5 years to 35 years and hence: (710-580)/(5-35) = −4.3 cm^3^/year). To reproduce their analysis and assess the effect of using the CNN-derived quality control estimates, we estimated gray and white matter volume in the subjects that had been scored for quality using our algorithm. Figure 4 shows gray matter volume as a function of age. Two conditions are compared: in one (Figure 4A) all of the subjects are included, while in the other only subjects that were passed by the CNN are included (Figure 4B, blue points). Depending on the threshold chosen, the effect of gray matter volume over age varies from −2.6 cm^3^/year (with no threshold) to −5.3 cm^3^/year (with Braindr rating > 0.9). A threshold of 0.7 of either Braindr or MRIQC results in an effect size around −4.3 cm^3^ per year, replicating the results of (Lebel and Beaulieu, 2011). A supplemental interactive version of this figure allows readers to threshold data points based on QC scores from the predicted labels of the CNN (called “Braindr ratings”), or on MRIQC XGBoost probabilities (called “MRIQC ratings”) is available at http://results.braindr.us. Thus, quality control has a substantial impact on estimates of brain development and allowing poor quality data into the statistical model can almost entirely obscure developmental changes in gray matter volume.

**Figure 4:**
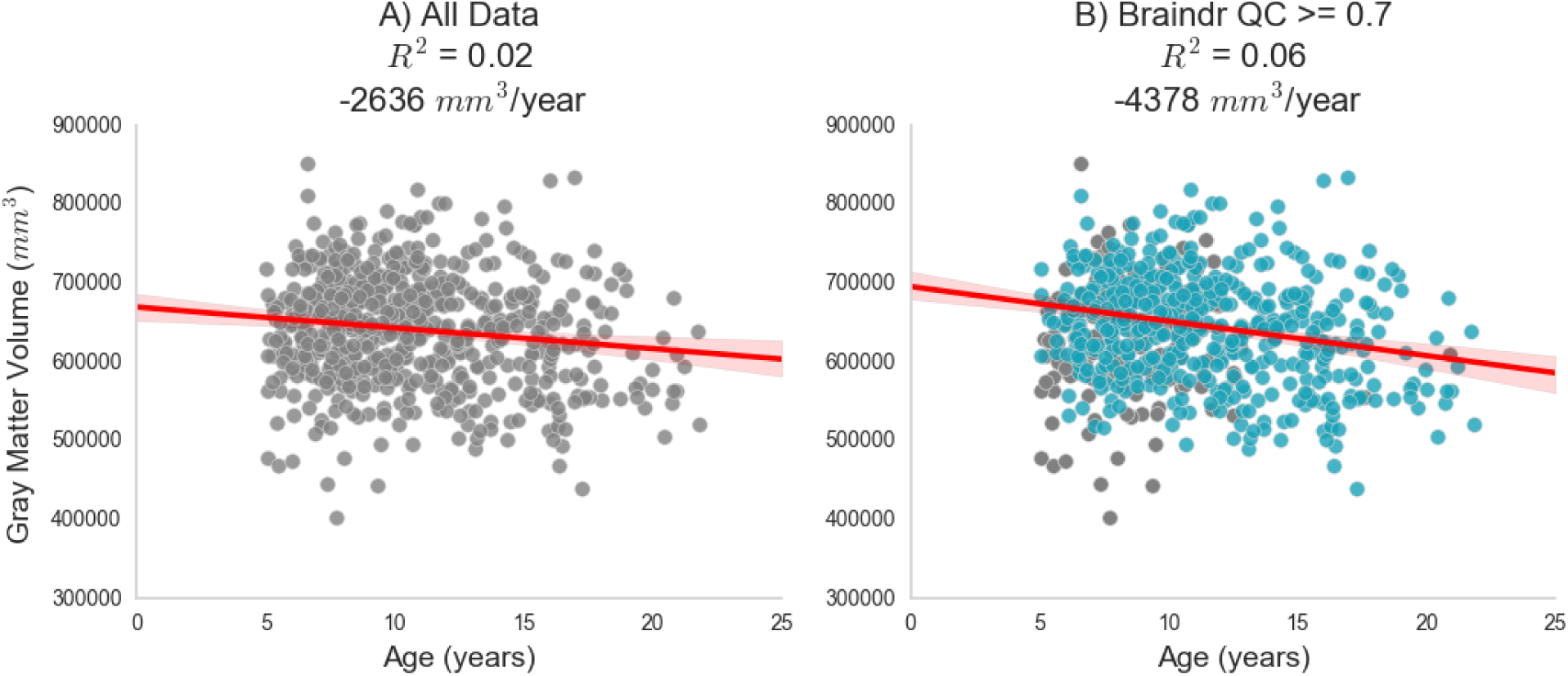
Impact of quality control on effect size estimates: Results of quality control on the inferred association between gray matter volume and age during development. Part A shows the relationship when all data is used in the ordinary least squares (OLS) model. Part B shows the new OLS model when data is thresholded by the deep learning model’s predicted braindr rating at 0.7. The effect size nearly doubles when QC scores are taken into account.

## Discussion

We have developed a system to scale expertise in neuroimaging to meet the demands of Big Data. The system uses citizen scientists to amplify an initially-small, expert-labeled dataset. Combined with deep learning (via CNNs), the system can then accurately perform image analysis tasks that require expertise, such as quality control (QC). We have validated our method against MRIQC, a specialized tool that was designed specifically for this use case based on knowledge of the physics underlying the signal generation process in T1-weighted images (Esteban et al., 2017). Unlike MRIQC, our method is able to generalize beyond quality control of T1-weighted images; any image-based binary classification task can be loaded onto the Braindr platform, and crowdsourced via the web. For this use-case, we demonstrated the importance of scaling QC expertise by showing how replication of a previously established results depends on a researcher’s decision on data quality. Lebel and colleagues (Lebel and Beaulieu, 2011) report changes in gray matter volume over development and we find that we only replicate these findings when using a stringent quality control threshold for the input data.

### The Internet and Web Applications for Collaboration

The internet and web browser technologies are not only crucial for scientific communication, but also for collaboration and distribution of work. This is particularly true in the age of large consortium efforts aimed at generating high-quality large data sets. Recent progress in citizen science projects for neuroscience research have proven extremely useful and popular, in part due to the ubiquity of the web browser. Large-scale citizen science projects, like EyeWire (Kim et al., 2014; Marx, 2013), and Mozak (Roskams and Popović, 2016), have enabled scientists working with high resolution microscopy data to map neuronal connections at the microscale, with help from over 100,000 citizen scientists. In MR imaging, web-based tools such as BrainBox (Heuer et al., 2016) and Mindcontrol (Keshavan et al., 2017) were built to facilitate the collaboration of neuroimaging experts in image segmentation and quality control. However, the task of inspecting each slice of a 3D image in either BrainBox or Mindcontrol takes a long time, and this complex task tends to lose potential citizen scientists who find it too difficult or time consuming. In general, crowdsourcing is most effective when a project is broken down into short, simple, well-defined “micro-tasks”, that can be completed in short bursts of work and are resilient to interruption (Cheng et al., 2015). In order to simplify the task for citizen scientists, we developed a web application called braindr, which reduces the time-consuming task of slice-by-slice 3D inspection to a quick binary choice made on a 2D slice. While we might worry that distilling a complex decision into a simple swipe on a smartphone might add noise, we demonstrated that a model could be constructed to accurately combine ratings from many citizen scientists to almost perfectly emulate those obtained from inspection by experts. Using braindr, citizen scientists amplified the initial expert-labelled dataset (200 3D images) to the entire dataset (> 700 3D images, > 3000 2D slices) in a few weeks. Because braindr is a lightweight web application, users could play it at any time and on any device, and this meant we were able to attract many users. On braindr, each slice received on average 20 ratings, and therefore each 3D brain (consisting of 5 slices) received on average 100 ratings. In short, by redesigning the way we interact with our data and presenting it in the web browser, we were able to get many more eyes on our data than would have been possible in a single research lab.

### Scaling expertise through interactions between experts, citizen scientists and machine learning

We found that an interaction between experts, citizen scientists, and machine learning results in scalable decision-making on brain MRI images. Recent advances in machine learning have vastly improved image classification(Krizhevsky et al., 2012), object detection(Girshick et al., 2014), and segmentation(Long et al., 2015) through the use of deep convolutional neural networks. In the biomedical domain, these networks have been trained to accurately diagnose eye disease (Lee et al., 2017a), diagnose skin cancer (Esteva et al., 2017), and breast cancer (Sahiner et al., 1996), to name a few applications. But these applications require a large and accurately labeled dataset. This presents an impediment for many scientific disciplines, where labeled data may be more scarce, or hard to come by, because it requires labor-intensive procedures. The approach presented here solves this fundamental bottleneck in the current application of modern machine learning approaches, and enables scientists to automate complex tasks that require substantial expertise.

A surprising finding that emerges from this work is that a deep learning algorithm can learn to match or even exceed the aggregated ratings that are used for training. This finding is likely to reflect the fact that algorithms are more reliable than humans, and when an algorithm is trained to match human accuracy, it has the added benefit of perfect reliability. For example even an expert might not provide the exact same ratings each time they see the same image, while an algorithm will. This is in line with findings from (Lee et al., 2017c), showing that the agreement between an algorithm and any one expert can be equivalent to agreement between any pair of experts. We have demonstrated that while an individual citizen scientists may not provide reliable results, by intelligently combining a crowd with machine learning, and keeping an expert in the loop to monitor results, decisions can be accurately scaled to meet the demands of Big Data.

### MRI Quality Control and Morphometrics over Development

The specific use-case that we focused on pertains to the importance of quality control in large-scale MRI data acquisitions. Recently, Ducharme and colleagues (Ducharme et al., 2016a) stressed the importance of quality control for studies of brain development in a large cohort of 954 subjects. They estimated cortical thickness on each point of a cortical surface and fit linear, quadratic and cubic models of thickness versus age at each vertex. Quality control was performed by visual inspection of the reconstructed cortical surface, and removing data that failed QC from the analysis. Without stringent quality control, the best fit models were more complex (quadratic/cubic), and with quality control the best fit models were linear. They found sex differences only at the occipital regions, which thinned faster in males. In the supplemental figure that accompanies Figure 4, we presented an interactive chart where users can similarly explore different ordinary least squares models (linear or quadratic) and also split by sex for the relationship between total gray matter volume, white matter volume, CSF volume, and total brain volume over age.

We chose to QC raw MRI data in this study, rather than the processed data because the quality of the raw MRI data affects the downstream cortical mesh generation, and many other computed metrics. A large body of research in automated QC of T1-weighted images exists, in part because of large open data sharing initiatives. In 2009, Mortamet and colleagues (Mortamet et al., 2009) developed a QC algorithm based on the background of magnitude images of the Alzheimer’s Disease Neuroimaging Initiative (ADNI) dataset, and reported a sensitivity and specificity of > 85%. In 2015, Shezad and colleagues (Shehzad et al., 2015) developed the Preprocessed Connectomes Project Quality Assessment Protocol (PCP-QAP) on the Autism Brain Imaging Data Exchange (ABIDE) and Consortium for Reproducibility and Reliability (CoRR) datasets. The PCP-QAP also included a Python library to easily compute metrics such as signal to noise ratio, contrast to noise ratio, entropy focus criterion, foreground-to-background energy ratio, voxel smoothness, and percentage of artifact voxels. Building on this work, the MRIQC package from Esteban and colleagues (Esteban et al., 2017) includes a comprehensive set of 64 image quality metrics, from which a classifier was trained to predict data quality of the ABIDE dataset for new, unseen sites with 76% accuracy.

Our strategy differed from that of the MRIQC classification study. In the Esteban 2017 study (Esteban et al., 2017), the authors labelled images that were “doubtful” in quality as a “pass” when training and evaluating their classifier. Our MRIQC classifier was trained and evaluated only on images that our raters very confidently passed or failed. Because quality control is subjective, we felt that it was acceptable for a “doubtful” image to be failed by the classifier. Since our classifier was trained on data acquired within a single site, and only on images that we were confident about, our MRIQC classifier achieved near perfect accuracy with an AUC of 0.99. On the other hand, our braindr CNN was trained as a regression (rather than a classification) on the full dataset, including the “doubtful” images (i.e those with ratings closer to 0.5), but was still evaluated as a classifier against data we were confident about. This also achieved near-perfect accuracy with an AUC of 0.99. Because both the MRIQC and braindr classifiers perform so well on data we are confident about, we contend that it is acceptable to let the classifier act as a “tie-breaker” for images that lie in the middle of the spectrum, for future acquisitions of the HBN dataset.

Quality control of large consortium datasets, and more generally, the scaling of expertise in neuroimaging, will become increasingly important as neuroscience moves towards data-driven discovery. Interdisciplinary collaboration between domain experts and computer scientists, and public outreach and engagement of citizen scientists can help realize the full potential of Big Data.

### Limitations

One limitation of this method is that there is an interpretability to speed tradeoff. Specialized QC tools were developed over many years, while this study was performed in a fraction of that time. Specialized QC tools are far more interpretable; for example, the coefficient of joint variation (CJV) metric from MRIQC is sensitive to the presence of head motion. CJV was one of the most important features of our MRIQC classifier, implying that our citizen scientists were primarily sensitive to motion artifacts. This conclusion is difficult to come to when interpreting the braindr CNN. Because we employed transfer learning, the features that were extracted were based on the ImageNet classification task, and it is unclear how these features related to MRI-specific artifacts. However, interpretability of deep learning is an ongoing active field of research (Chakraborty et al., 2017), and we may be able to fit more interpretable models in the future.

Compared to previous efforts to train models to predict quality ratings, such as MRIQC (Esteban et al., 2017), our AUC scores are very high. There are two main reasons for this. First, in the Esteban 2017 study (Esteban et al., 2017), the authors tried to predict the quality of scans from unseen sites, whereas in our study, we combined data across the two sites from which data had been made publicly available at the time we conducted this study. Second, even though our quality ratings on the 3D dataset were continuous scores (ranging from −5 to 5), we only evaluated the performance of our models on data that received an extremely high (4,5) or extremely low score (−4,−5) by the experts. This was because quality control is very subjective, and therefore, there is more variability on images that people are unsure about. An image that was failed with low confidence (−3 to −1) by one researcher could conceivably be passed with low confidence by another researcher (1 to 3). Most importantly, our study had enough data to exclude the images within this range of relative ambiguity in order to train our XGBoost model on both the braindr ratings and the MRIQC features. In studies with less data, such an approach might not be feasible.

Another limitation of this method was that our citizen scientists were primarily neuroscientists. The braindr application was advertised on Twitter (https://www.twitter.com) by the authors, whose social networks (on this platform) primarily consisted of neuroscientists. As the original tweet travelled outside our social network, we saw more citizen scientists without experience looking at brain images on the platform, but the number of ratings they contributed were not as high as those with neuroscience experience. We also saw that there was an overall tendency for all our users to incorrectly pass images. Future iterations of braindr will include a more informative tutorial and random checks with known images throughout the game to make sure our players are well informed and are performing well throughout the task. In this study, we were able to overcome this limitation because we had enough ratings to train the XGBoost algorithm to preferentially weight some user’s ratings over others.

### Future Directions

Citizen science platforms like the Zooniverse (Simpson et al., 2014) enable researchers to upload tasks and engage over 1 million citizen scientists. We plan to integrate braindr into a citizen science platform like Zooniverse. This would enable researchers to upload their own data to braindr, and give them access to a diverse group of citizen scientists, rather than only neuroscientists within their social network. We also plan to reuse the braindr interface for more complicated classification tasks in brain imaging. An example could be the classification of ICA components as signal or noise (Griffanti et al., 2017), or the evaluation of segmentation algorithms. Finally, incorporating braindr with existing open data initiatives, like OpenNeuro (Gorgolewski et al., 2017a), or existing neuroimaging platforms like LORIS (Das et al., 2012) would enable scientists to directly launch braindr tasks from these platforms, which would seamlessly incorporate human in the loop data analysis in neuroimaging research. More generally, the principles described here motivate platforms that integrate citizen science with deep learning for Big Data applications across the sciences.

## Methods

### The Healthy Brain Network Dataset

The first two releases of the Healthy Brain Network dataset were downloaded from http://fcon_1000.projects.nitrc.org/indi/cmi_healthy_brain_network/sharing_neuro.html. A web application for brain quality control, called Mindcontrol (Keshavan et al., 2017) was hosted at https://mindcontrol-hbn.herokuapp.com, which enabled users to view and rate 3D MRI images in the browser. There were 724 T1-weighted images. All procedures were approved by the University of Washington Institutional Review Board (IRB). Mindcontrol raters, who were all neuroimaging researchers with substantial experience in similar tasks, provided informed consent, including consent to publicly release these ratings. Mindcontrol raters were asked to pass or fail images after inspecting the full 3D volume, and provide a score of their confidence on a 5 point Likert scale, where 1 was the least confident and 5 was the most confident. Mindcontrol raters received a point for each new volume they rated, and a leaderboard on the homepage displayed rater rankings. The ratings of the top 4 expert raters (including the lead author) were used to create a gold-standard subset of the data.

### Gold-standard Selection

The gold-standard subset of the data was created by selecting images that were confidently passed or confidently failed (confidence equal or larger than 4) by the 4 expert raters. In order to measure reliability between expert raters, the ratings of the second, third, and fourth expert expert rater were recoded to a scale of −5 to 5 (where −5 is confidently failed, and 5 is confidently passed). An ROC analysis was performed against the binary ratings of the lead author on the commonly rated images, and the area under the curve (AUC) was computed for each pair. An average AUC, weighted by the number of commonly rated images between the pair, was 0.97, showing good agreement between expert raters. The resulting gold-standard dataset consisted of 200 images. Figure 5 shows example axial slices from the gold-standard dataset. The gold-standard dataset set contains 100 images that were failed by experts, and 100 images that were passed by experts.

**Figure 5:**
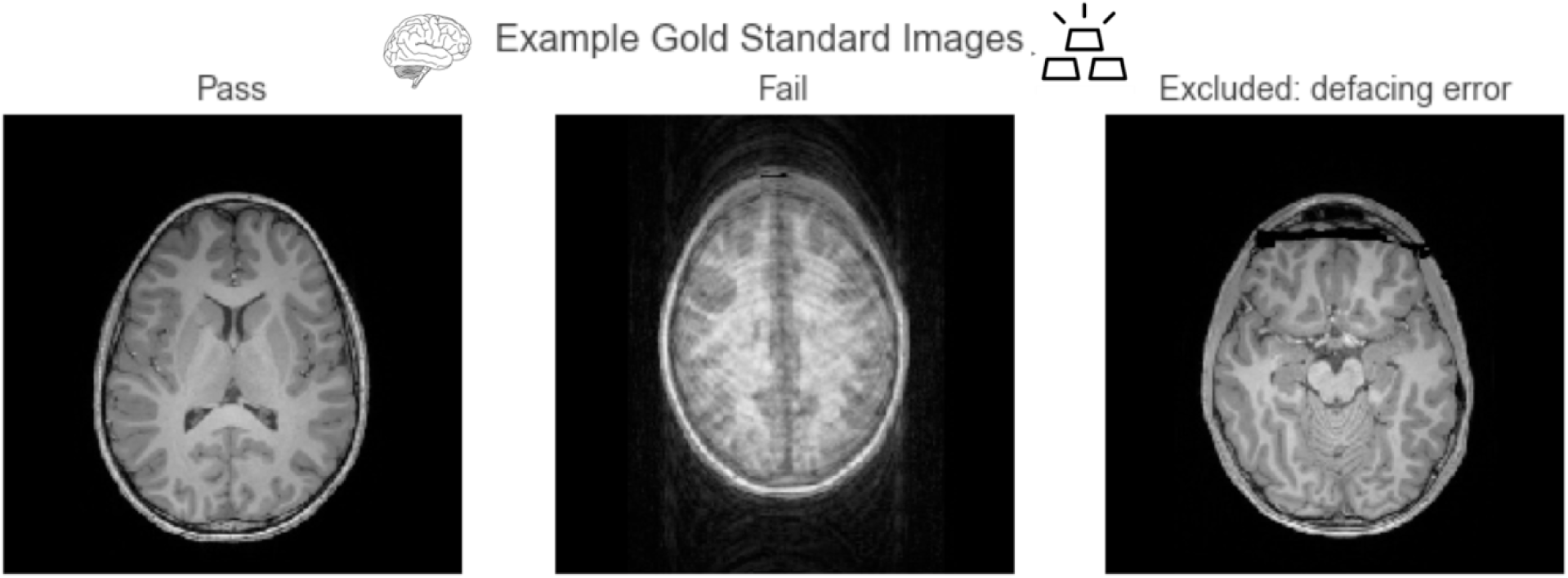
Example axial slices from the gold-standard dataset: Passed images show clear contrast between tissue types, and failed images primarily consisted of those with large motion artifacts. We excluded images that failed because of defacing errors from this analysis.

### Data Preparation

All images were then converted into a set of 2D axial slices using the NiBabel Python library (Brett et al., 2018) and uploaded to https://braindr.us. Two images of the 724 were corrupted, so the total image count became 722 images. Five slices, separated by 40 slices, were selected from each brain, where the first slice was one that had over 10,000 non-zero pixels. All slices were padded to 256×256 or 512×512 depending on original image size. One subject (sub-NDARVJ504DAA) had only 4 slices because the last slice did not meet the 10,000 pixel threshold. The total number of slices uploaded to https://braindr.us was 3609.

### The braindr web application

The braindr application was written in Javascript using the Vue.js (https://vuejs.org) framework. Google Firebase (https://firebase.google.com/) was used for the realtime database. The axial brain slices were hosted on Amazon S3 and served over the Amazon CloudFront content delivery network. Figure 6 shows the braindr interface, which presents to the user a 2D slice. On a touchscreen device (tablet or mobile phone), users can swipe right to pass or swipe left to fail the image. On a desktop, a user may click the “pass” or “fail” button or use the right or left arrow keys to classify the image. The user receives a point for each rating, unless they rate against the majority, where the majority is defined only for images with more than 5 ratings, and where the average rating is below 0.3 or above 0.7. The user receives a notification of the point they earned (or did not earn) for each image after each swipe. All users electronically signed a consent form as approved by the University of Washington IRB. Images were initially served randomly, and then images with fewer ratings were preferentially served.

**Figure 6:**
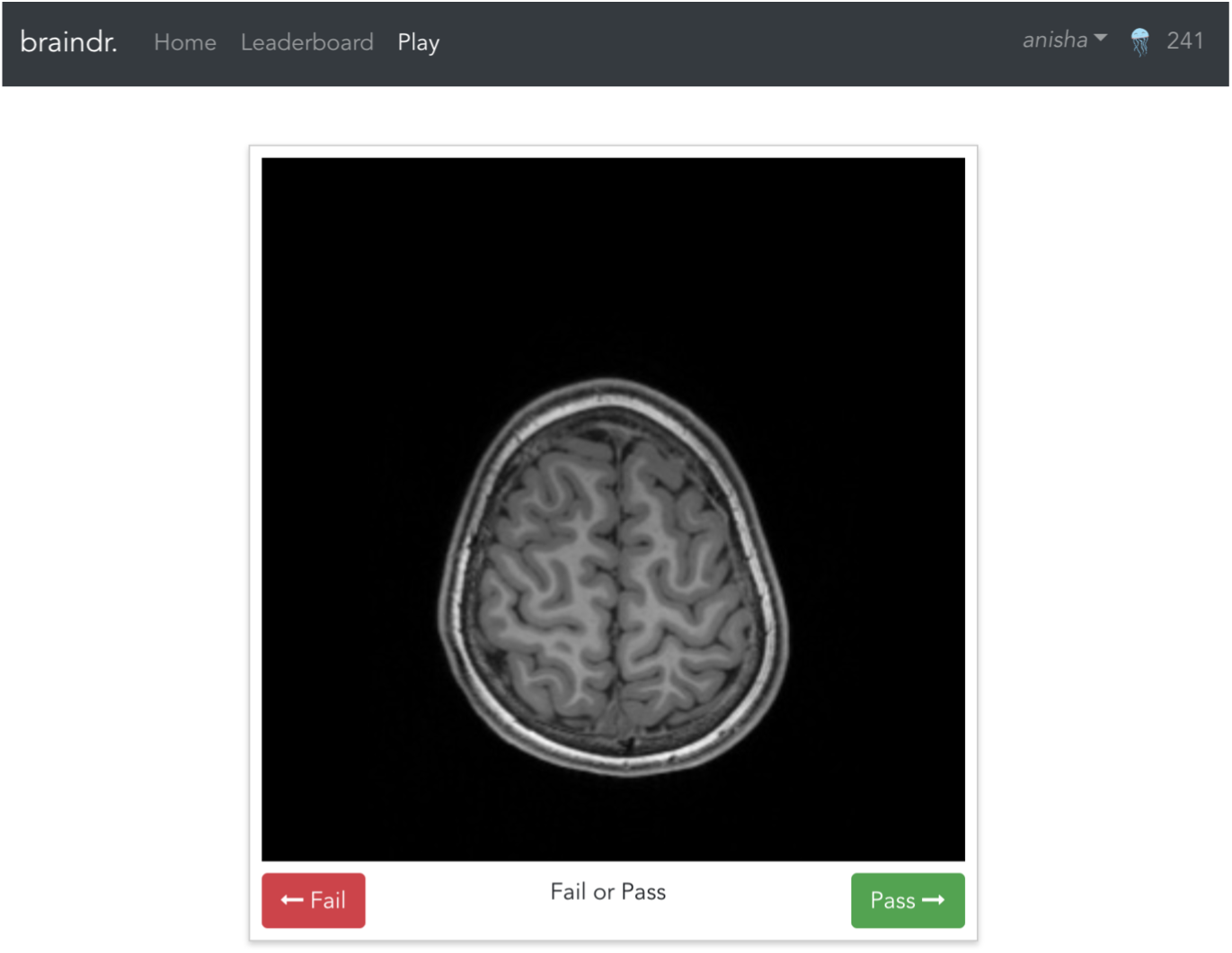
The braindr web interface: Braindr was hosted at https://braindr.us. Users may click pass or fail buttons, use arrow keys, or swipe on a touchscreen device to rate the image. The top right shows the user’s score.

### Braindr data collection

A total of 261 users submitted over 80,000 ratings. We selected the 25% of the users who rated the largest numbers of the gold-standard slices. This reduced the dataset to 65 users who submitted 68,314 total ratings, 18,940 of which were on the 1000 gold-standard slices. Figure 7 shows the distribution of average ratings and the distribution of number of ratings per slice on the gold-standard dataset.

**Figure 7:**
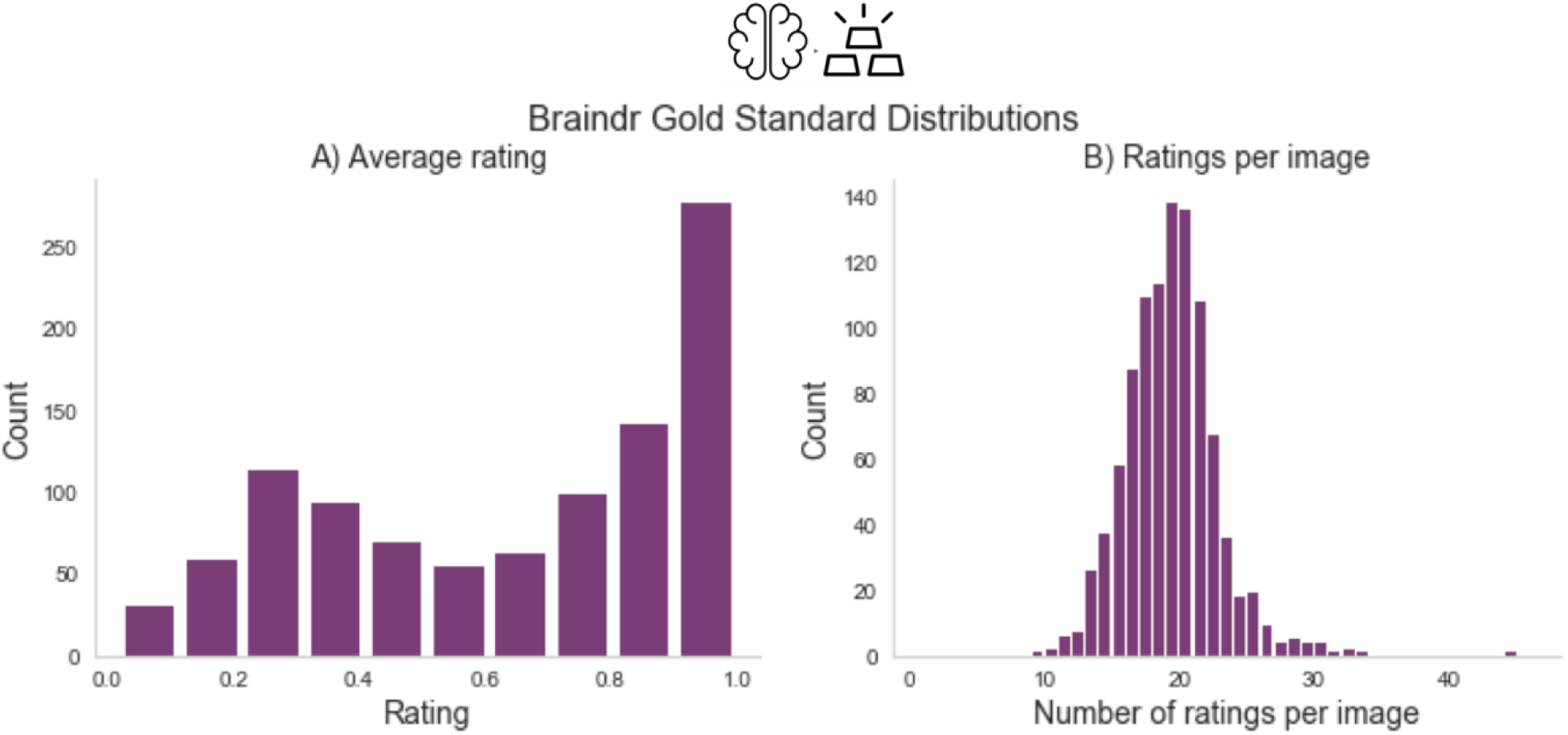
Braindr data distributions: Part A shows the distribution of average ratings for each slice on the gold-standard slices. Part B shows the number of ratings per slice, where on average each slice received 20 ratings.

### Rating aggregation with XGBoost

To aggregate citizen scientist ratings, we weighted citizen scientists based on how consistent their ratings were with the gold-standard. We trained an XGBoost classifier (Chen and Guestrin, 2016b) implemented in Python (http://xgboost.readthedocs.io/en/latest/python/python_intro.html) using the cross-validation functions from the scikit-learn Python library (Pedregosa et al., 2011). We used 600 estimators, and grid searched over a stratified 10-fold cross-validation within the training set to select the optimal maximum depth (2 vs 6) and learning rate (0.01, 0.1). The features of the model were the citizen scientists and each observation was a slice, with the entries in the design matrix set to be the average rating of a specific citizen scientist on a particular slice. We trained the classifier on splits of various sizes of the data to test the dependence on training size (see Figure 2A). We used the model trained with n=670 to extract the probability scores of the classifier on all 3609 slices in braindr (see Figure 2B). While equally weighting each citizen scientist’s ratings results in a bimodal distribution with a lower peak that is shifted up from zero (Figure 7A), the distribution of probability scores in Figure 2B more accurately matches our expectations of the data; a bimodal distribution with peaks at 0 and 1. Feature importances were extracted from the model and plotted in Figure 2C, and plotted against total number of gold-standard image ratings in Figure 2D.

### Deep learning to predict image QC label

Finally, a deep learning model was trained on the brain slices to predict the XGBoost probability score. All brain slices were resized to 256 by 256 pixels and converted to 3 color channels (RGB) to be compatible with the VGG16 input layer. The data was split into 80%-10%-10% training-validation-test sets. The data was split such that all slices belonging to the same subject were grouped together, so that any individual subject could be only in either training, validation or test. We loaded the VGG16 network that was pretrained with ImageNet weights (Simonyan and Zisserman, 2014) implemented in Keras (Chollet et al., 2015), removed the top layer, and ran inference on all the data. The output of the VGG16 inference was then used to train a small sequential neural network consisting of a dense layer with 256 nodes and a rectified linear unit activation function (ReLu), followed by a dropout layer set to drop 50% of the weights to prevent overfitting, and finally a single node output layer with sigmoid activation. The training of the final layer was run for 50 epochs and the best model on the validation set across the 50 epochs was saved. We ran this model 10 separate times, each time with a different random initialization seed, in order to measure the variability of our ROC AUC on the test set.

### Training the MRIQC model

MRIQC was run on all images in the HBN dataset. Rather than using the previously trained MRIQC classifier from Esteban and colleagues (Esteban et al., 2017), the extracted QC features were used to train another XGBoost classifier to predict gold-standard labels. Two thirds of the data was used to train the model, where a 2-fold cross-validation was used to optimize hyper parameters: learning rate = 0.001, 0.01, 0.1, number of estimators = 200, 600, and maximum depth = 2,6,8. An ROC analysis was run, and the computed area under the curve was 0.99.

### Gray matter volume vs age during development

Finally, to explore the relationship between gray matter volume and age over development as a function of QC threshold, gray matter volume was computed from running the Mindboggle software (Klein et al., 2017) on the entire dataset. Mindboggle combines the image segmentation output from Freesurfer (Fischl, 2012) with that of ANTS (Avants et al., 2011) to improve the accuracy of segmentation, labeling and volume shape features. Extremely low quality scans did not make it through the entire Mindboggle pipeline, and as a result the dataset size was reduced to 629 for this part of the analysis. The final QC score for the brain volumes was computed by taking the average of the predicted braindr rating from the deep learning model for all five slices. We ran an ordinary least squares (OLS) model on gray matter volume versus age on the data with and without QC thresholding, where the QC threshold was set at 0.7. Figure 4 shows the result of this analysis, which showed an effect size that nearly doubled and replicated previous findings when QC was performed on the data.

## Acknowledgements

This research was supported through a grant from the Gordon and Betty Moore Foundation and the Alfred P. Sloan Foundation to the University of Washington eScience Institute. A.K is also supported through a fellowship from the eScience Institute and the University of Washington Institute for Neuroengineering. We thank the NVIDIA corporation for supporting our research through their GPU seed grant. We’d like to acknowledge the following people for fruitful discussions and contributions to the project. Dylan Nielson, Satra Ghosh and Dave Kennedy for the inspiration for braindr. Greg Kiar, for contributing badges to the braindr application, and naming it. Chris Markiewicz, for discussions on application performance, and for application testing in the early stages. Katie Bottenhorn, Dave Kennedy, and Amanda Easson for quality controlling the gold-standard dataset. Jamie Hanson, for sharing the MRIQC metrics. Chris Madan, for application testing and for discussions regarding QC standards. Arno Klein and Lei Ai, for providing us the segmented images from the HBN dataset. Tal Yarkoni and Alejandro de la Vega, for organizing a “code rodeo” for neuroimagers in Austin, TX, where the idea for braindr was born. Finally, we’d like to thank all the citizen scientists who swiped on braindr - we are very grateful for your contributions!

## Code and Data Availability

The code for the braindr application can be found at https://doi.org/10.5281/zenodo.1208140. The brain slice data is hosted at https://osf.io/j5d4y/. The code for the analysis for this project, including all figures and the source code for the interactive version of this manuscript, can be found at https://github.com/akeshavan/braindr-results and https://github.com/akeshavan/braindr-analysis.

## References

Amr Ahmed, Kai Yu, Wei Xu, Yihong Gong, and Eric Xing. Training hierarchical feed-forward visual recognition models using transfer learning from pseudo-tasks. In European Conference on Computer Vision, pages 69–82. Springer, 2008. doi: 10.1007/978-3-540-88690-7_6.

Lindsay M Alexander, Jasmine Escalera, Lei Ai, Charissa Andreotti, Karina Febre, Alexander Mangone, Natan Vega-Potler, Nicolas Langer, Alexis Alexander, Meagan Kovacs, et al. An open resource for transdiagnostic research in pediatric mental health and learning disorders. Scientific data, 4:170181, 2017. doi: 10.1038/sdata.2017.181.

John Ashburner and Karl J. Friston. Unified segmentation. NeuroImage, 26(3):839–851, jul 2005. doi: 10.1016/j.neuroimage.2005.02.018. URL https://doi.org/10.1016%2Fj.neuroimage.2005.02.018.

Brian B. Avants, Nicholas J. Tustison, Gang Song, Philip A. Cook, Arno Klein, and James C. Gee. A reproducible evaluation of ANTs similarity metric performance in brain image registration. NeuroImage, 54(3):2033–2044, feb 2011. doi: 10.1016/j.neuroimage.2010.09.025. URL https://doi.org/10.1016%2Fj.neuroimage.2010.09.025.

NC Benson, OH Butt, R Datta, PD Radoeva, DH Brainard, and GK Aguirre. The retinotopic organization of striate cortex is well predicted by surface topology. Curr Biol, 22:2081–5, nov 2012. doi: 10.1016/j.cub.2012.09.014.

NC Benson, OH Butt, DH Brainard, and GK Aguirre. Correction of distortion in flattened representations of the cortical surface allows prediction of V1-V3 functional organization from anatomy. PLoS Comput Biol, 10:e1003538, Mar 2014. doi: 10.1371/journal.pcbi.1003538.

Matthew Brett, Michael Hanke, Chris Markiewicz, Marc-Alexandre Côté, Paul McCarthy, Satrajit Ghosh, Demian Wassermann, Stephan Gerhard, Yaroslav Halchenko, Eric Larson, Gregory R. Lee, Erik Kastman, Cindee M, Félix C. Morency, moloney, Ariel Rokem, Michiel Cottaar, Jarrod Millman, Ross Markello, jaeilepp, Chris Cheng, Alexandre Gramfort, Robert D Vincent, Jasper J.F. van den Bosch, Krish Subramaniam, Pradeep Reddy Raamana, Mathias Goncalves, Nolan Nichols, embaker, and Basile. nipy/nibabel: 2.3.0, jun 2018. URL https://doi.org/10.5281/zenodo.1287921.

BJ Casey, Tariq Cannonier, May I Conley, Alexandra O Cohen, Deanna M Barch, Mary M Heitzeg, Mary E Soules, Theresa Teslovich, Danielle V Dellarco, Hugh Garavan, et al. The adolescent brain cognitive development (ABCD) study: imaging acquisition across 21 sites. Developmental cognitive neuroscience, 2018.

M Catani and M Thiebautdeschotten. A diffusion tensor imaging tractography atlas for virtual in vivo dissections. Cortex, 44(8):1105–1132, sep 2008. doi: 10.1016/j.cortex.2008.05.004. URL https://doi.org/10.1016%2Fj.cortex.2008.05.004.

Supriyo Chakraborty, Richard Tomsett, Ramya Raghavendra, Daniel Harborne, Moustafa Alzantot, Federico Cerutti, Mani Srivastava, Alun Preece, Simon Julier, Raghuveer M Rao, et al. Interpretability of deep learning models: a survey of results. DAIS, 2017.

Tianqi Chen and Carlos Guestrin. Xgboost: A scalable tree boosting system. In Proceedings of the 22nd acm sigkdd international conference on knowledge discovery and data mining, pages 785–794. ACM, 2016a. doi: 10.1145/2939672.2939785.

Tianqi Chen and Carlos Guestrin. Xgboost: A scalable tree boosting system. In Proceedings of the 22nd acm sigkdd international conference on knowledge discovery and data mining, pages 785–794. ACM, 2016b.

Justin Cheng, Jaime Teevan, Shamsi T Iqbal, and Michael S Bernstein. Break it down: A comparison of macro-and microtasks. In Proceedings of the 33rd Annual ACM Conference on Human Factors in Computing Systems, pages 4061–4064. ACM, 2015.

François Chollet et al. Keras, 2015.

Özgün Çiçek, Ahmed Abdulkadir, Soeren S Lienkamp, Thomas Brox, and Olaf Ronneberger. 3D U-Net: learning dense volumetric segmentation from sparse annotation. In International Conference on Medical Image Computing and Computer-Assisted Intervention, pages 424–432. Springer, 2016. doi: 10.1007/978-3–319-46723-8_49.

Mehmet Ufuk Dalmiş, Geert Litjens, Katharina Holland, Arnaud Setio, Ritse Mann, Nico Karssemeijer, and Albert Gubern-Mérida. Using deep learning to segment breast and fibroglandular tissue in MRI volumes. Medical Physics, 44(2):533–546, feb 2017. doi: 10.1002/mp.12079. URL https://doi.org/10.1002%2Fmp.12079.

Samir Das, Alex P Zijdenbos, Dario Vins, Jonathan Harlap, and Alan C Evans. LORIS: a web-based data management system for multi-center studies. Frontiers in neuroinformatics, 5:37, 2012. doi: 10.3389/fninf.2011.00037.

Simon Ducharme, Matthew D Albaugh, Tuong-Vi Nguyen, James J Hudziak, JM Mateos-Pérez, Aurelie Labbe, Alan C Evans, Sherif Karama, Brain Development Cooperative Group, et al. Trajectories of cortical thickness maturation in normal brain development—The importance of quality control procedures. Neuroimage, 125:267–279, 2016a. doi: 10.1016/j.neuroimage.2015.10.010.

Simon Ducharme, Matthew D Albaugh, Tuong-Vi Nguyen, James J Hudziak, JM Mateos-Pérez, Aurelie Labbe, Alan C Evans, Sherif Karama, Brain Development Cooperative Group, et al. Trajectories of cortical thickness maturation in normal brain development—The importance of quality control procedures. Neuroimage, 125:267–279, 2016b. doi: 10.1016/j.neuroimage.2015.10.010.

Oscar Esteban, Daniel Birman, Marie Schaer, Oluwasanmi O Koyejo, Russell A Poldrack, and Krzysztof J Gorgolewski. MRIQC: Advancing the automatic prediction of image quality in MRI from unseen sites. PloS one, 12(9):e0184661, 2017. doi: 10.1371/journal.pone.0184661.

Andre Esteva, Brett Kuprel, Roberto A Novoa, Justin Ko, Susan M Swetter, Helen M Blau, and Sebastian Thrun. Dermatologist-level classification of skin cancer with deep neural networks. Nature, 542(7639): 115, 2017. doi: 10.1038/nature21056.

Jianqing Fan, Fang Han, and Han Liu. Challenges of Big Data analysis. National Science Review, 1(2): 293–314, feb 2014. doi: 10.1093/nsr/nwt032. URL https://doi.org/10.1093%2Fnsr%2Fnwt032.

Adam R Ferguson, Jessica L Nielson, Melissa H Cragin, Anita E Bandrowski, and Maryann E Martone. Big data from small data: data-sharing in the ‘long tail’ of neuroscience. Nature Neuroscience, 17(11): 1442–1447, nov 2014. doi: 10.1038/nn.3838. URL https://doi.org/10.1038%2Fnn.3838.

Bruce Fischl. FreeSurfer. Neuroimage, 62(2):774–781, 2012. doi: 10.1016/j.neuroimage.2012.01.021.

Ross Girshick, Jeff Donahue, Trevor Darrell, and Jitendra Malik. Rich feature hierarchies for accurate object detection and semantic segmentation. In Proceedings of the IEEE conference on computer vision and pattern recognition, pages 580–587, 2014. doi: 10.1109/CVPR.2014.81. URL https://arxiv.org/abs/1311.2524.

Matthew F Glasser, Stephen M Smith, Daniel S Marcus, Jesper LR Andersson, Edward J Auerbach, Timothy EJ Behrens, Timothy S Coalson, Michael P Harms, Mark Jenkinson, Steen Moeller, et al. The human connectome project’s neuroimaging approach. Nature Neuroscience, 19(9):1175, 2016.

K Gorgolewski, Oscar Esteban, Gunnar Schaefer, B Wandell, and R Poldrack. OpenNeuro—a free online platform for sharing and analysis of neuroimaging data. Organization for Human Brain Mapping. Vancouver, Canada, page 1677, 2017a.

Krzysztof J Gorgolewski, Tibor Auer, Vince D Calhoun, R Cameron Craddock, Samir Das, Eugene P Duff, Guillaume Flandin, Satrajit S Ghosh, Tristan Glatard, Yaroslav O Halchenko, et al. The brain imaging data structure, a format for organizing and describing outputs of neuroimaging experiments. Scientific Data, 3:160044, 2016.

Krzysztof J Gorgolewski, Fidel Alfaro-Almagro, Tibor Auer, Pierre Bellec, Mihai Capotă, M Mallar Chakravarty, Nathan W Churchill, Alexander Li Cohen, R Cameron Craddock, Gabriel A Devenyi, et al. BIDS apps: Improving ease of use, accessibility, and reproducibility of neuroimaging data analysis methods. PLoS computational biology, 13(3):e1005209, 2017b.

Ludovica Griffanti, Gwenaëlle Douaud, Janine Bijsterbosch, Stefania Evangelisti, Fidel Alfaro-Almagro, Matthew F Glasser, Eugene P Duff, Sean Fitzgibbon, Robert Westphal, Davide Carone, et al. Hand classification of fMRI ICA noise components. Neuroimage, 154:188–205, 2017. doi: 10.1016/j.neuroimage. 2016.12.036.

Katja Heuer, Satrajit Ghosh, Amy Robinson Sterling, and Roberto Toro. Open neuroimaging laboratory. Research Ideas and Outcomes, 2:e9113, 2016. doi: 10.3897/rio.2.e9113.

Kesshi M. Jordan, Bagrat Amirbekian, Anisha Keshavan, and Roland G. Henry. Cluster Confidence Index: A Streamline-Wise Pathway Reproducibility Metric for Diffusion-Weighted MRI Tractography. Journal of Neuroimaging, 28(1):64–69, sep 2017a. doi: 10.1111/jon.12467. URL https://doi.org/10.1111%2Fjon.12467.

Kesshi Marin Jordan, Anisha Keshavan, Eduardo Caverzasi, Joseph Osorio, Nico Papinutto, Bagrat Amirbekian, Mitchel S. Berger, and Roland G. Henry. Investigating The Functional Consequence Of White Matter Damage: An Automatic Pipeline To Create Longitudinal Disconnection Tractograms. may 2017b. doi: 10.1101/140137. URL https://doi.org/10.1101%2F140137.

Anisha Keshavan, Esha Datta, Ian M McDonough, Christopher R Madan, Kesshi Jordan, and Roland G Henry. Mindcontrol: A web application for brain segmentation quality control. NeuroImage, 2017. doi: 10.1016/j.neuroimage.2017.03.055.

Jinseop S Kim, Matthew J Greene, Aleksandar Zlateski, Kisuk Lee, Mark Richardson, Srinivas C Turaga, Michael Purcaro, Matthew Balkam, Amy Robinson, Bardia F Behabadi, et al. Space-time wiring specificity supports direction selectivity in the retina. Nature, 509(7500):331, 2014. doi: 10.1038/nature13240.

Arno Klein, Satrajit S Ghosh, Forrest S Bao, Joachim Giard, Yrjö Häme, Eliezer Stavsky, Noah Lee, Brian Rossa, Martin Reuter, Elias Chaibub Neto, et al. Mindboggling morphometry of human brains. PLoS computational biology, 13(2):e1005350, 2017. doi: 10.1371/journal.pcbi.1005350.

Alex Krizhevsky, Ilya Sutskever, and Geoffrey E Hinton. Imagenet classification with deep convolutional neural networks. In Advances in neural information processing systems, pages 1097–1105, 2012. doi: 10.1145/3065386.

Catherine Lebel and Christian Beaulieu. Longitudinal development of human brain wiring continues from childhood into adulthood. Journal of Neuroscience, 31(30):10937–10947, 2011. doi: 10.1523/JNEUROSCI.5302–10.2011.

Cecilia S Lee, Doug M Baughman, and Aaron Y Lee. Deep learning is effective for classifying normal versus age-related macular degeneration OCT images. Ophthalmology Retina, 1(4):322–327, 2017a. doi: 10.1016/j.oret.2016.12.009.

Cecilia S. Lee, Ariel J. Tyring, Nicolaas P. Deruyter, Yue Wu, Ariel Rokem, and Aaron Y. Lee. Deep-learning based automated segmentation of macular edema in optical coherence tomography. Biomedical Optics Express, 8(7):3440, jun 2017b. doi: 10.1364/boe.8.003440. URL https://doi.org/10.1364%2Fboe.8.003440.

Cecilia S Lee, Ariel J Tyring, Nicolaas P Deruyter, Yue Wu, Ariel Rokem, and Aaron Y Lee. Deep-learning based, automated segmentation of macular edema in optical coherence tomography. Biomedical optics express, 8(7):3440–3448, 2017c. doi: 10.1364/BOE.8.003440.

Jonathan Long, Evan Shelhamer, and Trevor Darrell. Fully convolutional networks for semantic segmentation. In Proceedings of the IEEE conference on computer vision and pattern recognition, pages 3431–3440, 2015. doi: 10.1109/TPAMI.2016.2572683.

Vivien Marx. Neuroscience waves to the crowd, 2013.

Bénédicte Mortamet, Matt A Bernstein, Clifford R Jack, Jeffrey L Gunter, Chadwick Ward, Paula J Britson, Reto Meuli, Jean-Philippe Thiran, and Gunnar Krueger. Automatic quality assessment in structural brain magnetic resonance imaging. Magnetic resonance in medicine, 62(2):365–372, 2009. doi: 10.1002/mrm.21992.

Sinno Jialin Pan and Qiang Yang. A survey on transfer learning. IEEE Transactions on knowledge and data engineering, 22(10):1345–1359, 2010. doi: 10.1109/TKDE.2009.191.

Brian Patenaude, Stephen M. Smith, David N. Kennedy, and Mark Jenkinson. A Bayesian model of shape and appearance for subcortical brain segmentation. NeuroImage, 56(3):907–922, jun 2011. doi: 10.1016/j.neuroimage.2011.02.046. URL https://doi.org/10.1016%2Fj.neuroimage.2011.02.046.

Fabian Pedregosa, Gaël Varoquaux, Alexandre Gramfort, Vincent Michel, Bertrand Thirion, Olivier Grisel, Mathieu Blondel, Peter Prettenhofer, Ron Weiss, Vincent Dubourg, et al. Scikit-learn: Machine learning in Python. Journal of machine learning research, 12(Oct):2825–2830, 2011.

Russell A Poldrack and Krzysztof J Gorgolewski. Making big data open: data sharing in neuroimaging. Nature Neuroscience, 17(11):1510–1517, oct 2014. doi: 10.1038/nn.3818. URL https://doi.org/10.1038%2Fnn.3818.

Olaf Ronneberger, Philipp Fischer, and Thomas Brox. U-net: Convolutional networks for biomedical image segmentation. In *International Conference on Medical image computing and computer-assisted intervention*, pages 234–241. Springer, 2015. doi: 10.1007/978-3-319-24574-4_28.

Jane Roskams and Zoran Popović. Power to the people: Addressing big data challenges in neuroscience by creating a new cadre of citizen neuroscientists. Neuron, 92(3):658–664, 2016. doi: 10.1016/j.neuron.2016.10.045.

Olga Russakovsky, Jia Deng, Hao Su, Jonathan Krause, Sanjeev Satheesh, Sean Ma, Zhiheng Huang, Andrej Karpathy, Aditya Khosla, Michael Bernstein, Alexander C. Berg, and Li Fei-Fei. ImageNet Large Scale Visual Recognition Challenge. International Journal of Computer Vision (IJCV), 115(3):211–252, 2015. doi: 10.1007/s11263-015-0816-y.

Berkman Sahiner, Heang-Ping Chan, Nicholas Petrick, Datong Wei, Mark A Helvie, Dorit D Adler, and Mitchell M Goodsitt. Classification of mass and normal breast tissue: a convolution neural network classifier with spatial domain and texture images. IEEE transactions on Medical Imaging, 15(5):598–610, 1996. doi: 10.1109/42.538937.

Takaya Saito and Marc Rehmsmeier. The precision-recall plot is more informative than the ROC plot when evaluating binary classifiers on imbalanced datasets. PloS one, 10(3):e0118432, 2015. doi: 10.1371/journal.pone.0118432.

Zarrar Shehzad, Steven Giavasis, Qingyang Li, Yassine Benhajali, Chaogan Yan, Zhen Yang, Michael Milham, Pierre Bellec, and Cameron Craddock. The Preprocessed Connectomes Project Quality Assessment Protocol: A resource for measuring the quality of MRI data. Frontiers in Neuroscience, (47), 2015. ISSN 1662-453X. doi: 10.3389/conf.fnins.2015.91.00047. URL http://www.frontiersin.org/10.3389/conf.fnins.2015.91.00047/event_abstract.

Karen Simonyan and Andrew Zisserman. Very deep convolutional networks for large-scale image recognition. arXiv preprint arXiv:1409.1556, 2014. URL https://arxiv.org/abs/1409.1556e.

Robert Simpson, Kevin R Page, and David De Roure. Zooniverse: observing the world’s largest citizen science platform. In Proceedings of the 23rd international conference on world wide web, pages 1049–1054. ACM, 2014. doi: 10.1145/2567948.2579215.

John D. Van Horn and Arthur W. Toga. Human neuroimaging as a “Big Data” science. Brain Imaging and Behavior, 8(2):323–331, oct 2013. doi: 10.1007/s11682-013-9255-y. URL https://doi.org/10.1007%2Fs11682-013-9255-y.

Annegreet Van Opbroek, M Arfan Ikram, Meike W Vernooij, and Marleen De Bruijne. Transfer learning improves supervised image segmentation across imaging protocols. IEEE transactions on medical imaging, 34(5):1018–1030, 2015. doi: 10.1109/TMI.2014.2366792.

BA Wandell and J Winawer. Imaging retinotopic maps in the human brain. Vision Res, 51:718–37, Apr 2011. doi: 10.1016/j.visres.2010.08.004.

J Winawer and N Witthoft. Identification of the ventral occipital visual field maps in the human brain. F1000Res, 6:1526, 2017. doi: 10.12688/f1000research.12364.1.

Jason D Yeatman, Robert F Dougherty, Nathaniel J Myall, Brian A Wandell, and Heidi M Feldman. Tract profiles of white matter properties: automating fiber-tract quantification. PloS one, 7(11):e49790, 2012.

